# Formin tails act as a switch, inhibiting or enhancing processive actin elongation

**DOI:** 10.1101/2023.09.06.556579

**Authors:** Kathryn Bremer, Carolyn Wu, Aanand Patel, Kevin He, Alex Grunfeld, Guillaume Chanfreau, Margot Quinlan

## Abstract

Formins are large, multidomain proteins that nucleate new actin filaments and accelerate elongation through a processive interaction with the barbed end of filaments. Their actin assembly activity is generally attributed to their eponymous Formin Homology (FH) 1 and 2 domains; however, evidence is mounting that regions outside of the FH1FH2 stretch also tune actin assembly. Here, we explore the underlying contributions of the tail domain, which spans the sequence between the FH2 domain and the C-terminus of formins. Tails vary in length from ∼0 to >200 residues and contain a number of recognizable motifs. The most well-studied motif is the ∼15 residue long diaphanous autoregulatory domain (DAD). The DAD mediates all or nothing regulation of actin assembly through an intramolecular interaction with the diaphanous inhibitory domain (DID) in the N-terminal half of the protein. Multiple reports demonstrate that the tail can enhance both nucleation and processivity. In this study, we provide a high-resolution view of the alternative splicing encompassing the tail in the <u>F</u>ormin <u>Ho</u>mology <u>D</u>omain-(Fhod) family of formins during development. While four distinct tails are predicted, we found significant levels of only two of these. We characterized the biochemical effects of the different tails. Surprisingly, the two highly expressed Fhod-tails inhibit processive elongation and diminish the nucleation and elongation rates, while a third supports activity. These findings demonstrate a new mechanism of modulating actin assembly by formins and support the model that splice variants are specialized to build distinct actin structures during development.

## Introduction

Formins are a highly conserved family of proteins that nucleate actin and modify filament growth by remaining processively associated with the fast-growing barbed end of actin filaments (1–4). They are defined by their homodimeric formin homology (FH)-2 domains and proline-rich FH1 domains. The FH2 domain dimerizes to form a donut-shaped structure that is sufficient for nucleation and processive binding at the barbed end of elongating filaments (5–7). The proline-rich FH1 domain recruits profilin-bound actin monomers and delivers them to the FH2-bound barbed end, accelerating the actin assembly rate (8, 9). In addition to the FH1/2 domains, most formins contain a loosely defined tail domain between the FH2 domain and the C-terminus. The tail often contains a diaphanous autoregulatory domain (DAD), which binds to an N-terminal diaphanous inhibitory domain (DID) to inhibit formin activity (10, 11). The tail also contributes to nucleation and elongation (12, 13).

The tail length varies greatly across formins, ranging from absent to >200 residues. By testing several truncations of the formin Cappuccino (Capu) and chimeras of tails from various formins added to the Capu-FH1FH2 domain, we previously found that the tail strongly influences processivity but not the elongation rate (13). The dissociation rate of Capu from the barbed end was over two orders of magnitude higher when the ∼30-residue tail of Capu was deleted. Processivity could be improved by replacing the Capu-tail with tails from the highly processive formins, mDia1 and mDia2. The dissociation rate loosely correlated with the pI of the tail. Strikingly, close to wild-type processivity was recovered when we scrambled the order of the Capu-tail residues. These observations led to a model in which the tail domain makes non-specific electrostatic contacts with the sides of growing filaments to enhance processivity, consistent with the idea that the pI may be a useful predictor of processivity.

The *Drosophila melanogaster* <u>F</u>ormin <u>Ho</u>mology 2 <u>D</u>omain family protein, referred to here as Fhod (the gene name is annotated as *fhos* or *knittrig (14)*), plays many roles in development, including myofibril assembly, tracheal development, and programmed autophagic cell death in salivary glands (15–17). In adults, Fhod plays a maintenance role in muscle cells and contributes to macrophage motility and the immune response (15, 16, 18). Fhod is alternatively spliced to produce as many as nine isoforms that differ by their N-termini and their tails but all contain identical FH1 and FH2 domains (14). Functional analysis demonstrates critical roles for at least two different N-termini (15, 16). However, there is no known role for the four Fhod tails that are generated by alternative splicing. We previously reported that the C-terminal half of one of these Fhod isoforms, Fhod-A, nucleates actin filaments and binds barbed ends, but is only weakly processive (19). To better understand the role of Fhod and the role of the formin tail in actin assembly, we exploited the natural variation of Fhod transcripts, comparing actin assembly activities of Fhod-FH1FH2 with different tails. In contrast to what we previously observed for Fhod-A, we found that a shorter tail (Fhod-B) supports processivity. In contrast to what we previously observed for Capu, we found that longer Fhod tails suppress processivity and decrease the nucleation and elongation rates. Nucleation and elongation rates correlate with tail length in Fhod, whereas processivity is impaired by a specific nine residues. Thus, formin tails can be highly specialized and even within a single gene, confer distinct actin assembly properties.

## Results

### Oxford Nanopore sequencing provides a high resolution view of Fhod variants expressed differentially during fly development

The *Drosophila* formin Fhod provides a powerful model to study contributions to actin assembly by the domains outside of the FH1 and FH2 domains. Only one *fhos* gene, which encodes the protein Fhod is found in the fly genome, but alternative splicing results in at least nine predicted protein products, which vary by their N-termini and C-terminal tails while retaining identical FH1 and FH2 domains (Fig. 1A).

**Figure 1.**
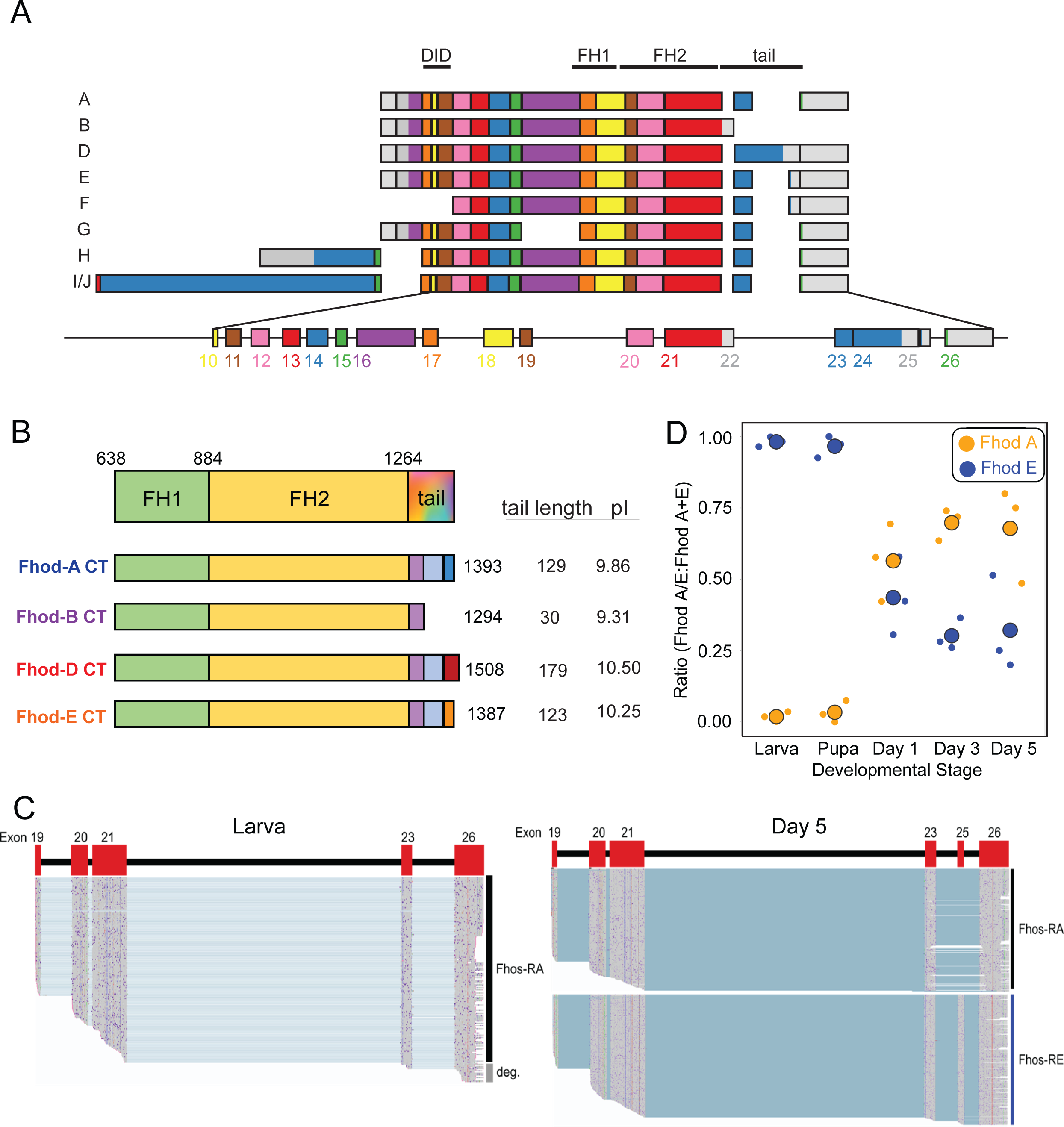
Fhod variants with distinct tails are differentially expressed during development. A) Alternative splicing of *fhos*, leads to nine putative variants. B) Four distinct tails are possible (A,B,D,E). They have the same Formin homology-1 (FH1) and FH2 domains. FH1 green; FH2 yellow; within the tail, colors represent exons. The tails vary in length and theoretical pI. C) Oxford Nanopore sequencing – samples of reads mapped to the Fhod gene for the larval stage and Day 5. Data from all five stages are shown in Supporting Information (Fig. S1). D) Ratio of reads of Fhod-A (blue) relative to reads of Fhod-E (yellow) for each developmental stage. Plot generated using SuperPlotsOfData (30). Small circles represent reads from independent samples. The large circles are means (n = 3 in all cases). Fhod-A is present at all stages while Fhod-E is not-detected in larva and pupa but read numbers increase with age post-eclosion.

We focused on the *fhos* isoforms that differ by their tails and asked which of the variants are expressed as a function of development. Note that there are two groups of isoforms that each share a common tail. For simplicity, we refer to the tail shared by isoforms A, G, H, I, and J as the Fhod-A tail and the tail shared by isoforms E and F as the Fhod-E tail (Fig.1A). The Fhod-B and Fhod-D tails are not shared by other isoforms.

To study expression patterns, we isolated total RNA from whole larvae, pupae, and days 1, 3, and 5 post-eclosion. We enriched these samples for Fhod, from the FH2 domain to the poly-A tail, using end-point RT-PCR (primer sequences are included in Methods). We then used Nanopore amplicon sequencing to determine which isoforms were expressed at different developmental stages. We detected Fhod-A in all developmental stages tested, while the Fhod-E tail was only observed in eclosed flies (Figure 1C, D). Data from all five developmental stages are shown in Supporting Information (Fig. S1). While Fhod-A levels decrease relative to Fhod-E, it is still expressed in adult flies as shown by the abundance of Fhod-A reads. Fhod-B and Fhod-D were not detected using this approach. This suggests that Fhod-B and Fhod-D may not be expressed or, if they are, they are possibly expressed in specific tissues and at levels too low to be detected in samples from whole animals.

Interestingly, we detected an extended species of exon 19, which terminated within the following intron (Fig. S2A). The FH2 domain spans exons 19, 20 and 21. This species would encode only a small portion of the FH2 domain. An additional species accumulated in adult flies that terminated at exon 21, which could be Fhod-B; however, it does not contain the distinct Fhod-B 3’UTR (Fig. S2B) although we did detect the Fhod-B 3’-UTR a few times (Fig. S2C). To our knowledge, neither of these transcripts has been previously detected.

In conclusion, the Nanopore sequencing data provided a high resolution picture of the expression of the different Fhod isoforms during development. Because of their distinct developmental expression patterns and apparently high levels of expression, we chose to focus on Fhod-A and Fhod-E. We included Fhod-B in our biochemical analysis because, when translated, it is effectively a truncation of the other three tails.

### Fhod-B is a processive elongation factor

In order to characterize the biochemical properties of the different Fhod tails, we purified the C-terminal halves of the two most highly expressed isoforms (A and E), which includes their common FH1 and FH2 domains and their distinct C-terminal tails (Fig. 1A). The tails of isoforms A, B, and E share the same first 30 residues. Fhod-B terminates at this point, whereas Fhod-A and Fhod-E have longer tails, consisting of 75 residues that are shared between the two followed by a short sequence unique to each (24 residues long for Fhod-A and 18 residues long for Fhod-E; Fig. 1B). Using TIRF microscopy, we did not detect obvious differences in elongation rate or fluorescence intensity when actin filaments were grown in the presence of 0.5 nM Fhod-A or Fhod-E, compared to actin alone (Fig. 2A), suggesting that the formins have short run lengths. (Formin-bound filaments appear dimmer because the FH1 domains recruit profilin-bound actin, and profilin has a weaker affinity for actin that is fluorescently labeled on cysteine (8).) These data are consistent with our previous study of Fhod-A, which has a characteristic run length of ∼2 μm, which is too short to reliably measure in typical TIRF microscopy assays (19). Due to their short run lengths, we could not determine if Fhod-A or Fhod-E alters the elongation rate when bound to barbed ends.

**Figure 2.**
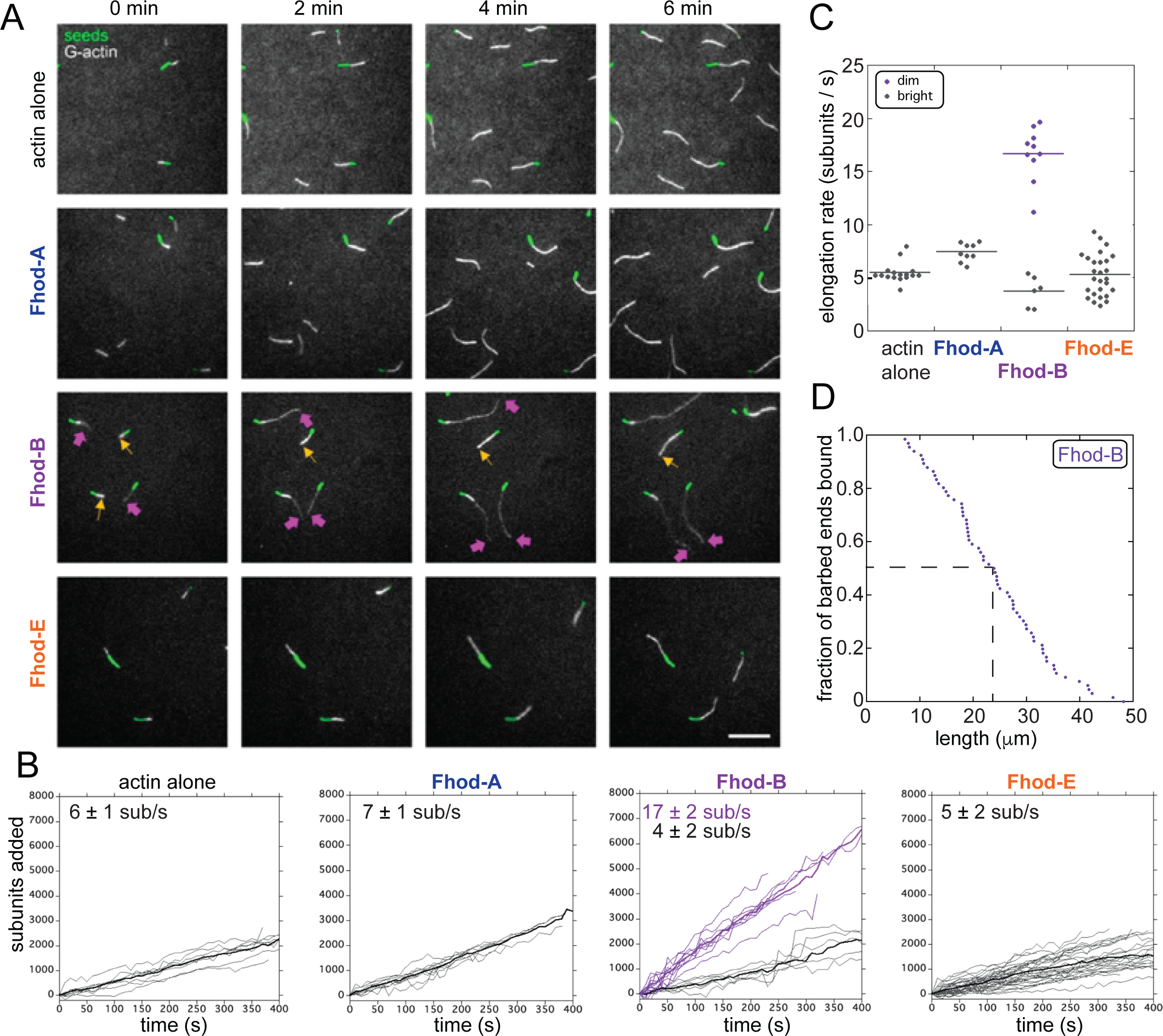
Fhod-B is a processive elongator. A) Direct observation of barbed-end elongation by TIRF microscopy with 0.5 nM seeds (1% biotinylated, labeled with Alexa Fluor 647-phalloidin; green), 1 μM G-actin (20% Oregon green-labeled; white), 5 μM Chic, ± 0.5 nM indicated Fhod construct. Yellow arrows denote the barbed-ends of bright, slow-growing filaments (no Fhod bound). Magenta arrows denote the barbed-ends of dim, fast-growing filaments (Fhod bound). *Scale bar,* 10 μm. C-F) Quantification of elongation from (B). Fine grey traces denote bright, slow-growing filaments. Fine purple traces denote dim, fast-growing filaments (Fhod bound). Thick grey and purple traces denote means of the bright and dim growing populations, respectively. Elongation rates at the top of each plot are the mean ± standard deviation from ≥ 2 flow chambers for each condition (*n* = 16, actin alone; *n* = 9, Fhod-A; *n* = 10 [dim] and *n* = 6 [bright], Fhod-B; *n* = 27, Fhod-E). C) Comparison of elongation rates for Fhod isoforms. Lines indicate the means. D) Measurement of Fhod-B processivity shown as an empirical cumulative distribution function (CDF) plot. Data points are from three experimental replicates (n = 66 filaments). The dashed line indicates the median of the combined samples (24 μm). The mean +/-standard deviation of the three independent samples is 24.1 +/-0.6 μm.

In contrast to Fhod-A and Fhod-E, growing F-actin seeds in the presence of 0.5 nM Fhod-B resulted in bright and dim filaments, where the dim filaments grew faster than their bright counterparts (17 ± 2 vs 4 ± 1 subunits/sec, respectively; Fig. 2A-C). We thus conclude that Fhod-B remains processively associated with the barbed end of filaments and accelerates elongation. We estimated the characteristic run length of Fhod-B by measuring the length distribution of actin filaments in this elongation assay after 5 minutes (Fig. 2D). The distribution of (dim) filament lengths indicates a median run length of 24 µm (n=66). The data are not well fit by an exponential, in part because we cannot detect the shortest filaments. In addition, this value is likely to be an underestimate of the characteristic run length because we are unable to measure filaments longer than 50 μm or that grow for longer than 5 minutes, with this assay.

Thus, we have identified a *Drosophila* Fhod isoform, Fhod-B, which elongates filaments with at least 10-fold increased processivity compared to the previously characterized isoform Fhod-A and the newly characterized isoform Fhod-E. The amino acid sequences of these isoforms only differ within their tail region, where Fhod-B possesses the shortest tail. Therefore, our data show that there is nothing inherent to Fhod-FH1FH2 that prevents processive elongation. Instead, the data indicate that Fhod processivity switches depending on its tail and that processivity is markedly decreased or even inhibited by certain tails.

### Residues near the end of the Fhod-A tail inhibit processivity

We sought to determine the basis of processivity loss, given that Fhod-A and Fhod-E include the sequence of Fhod-B. To address this, we first asked whether each tail adopts any secondary structure using circular dichroism (CD) spectroscopy. The spectra for all three isoforms were consistent with random coil, leading us to conclude that all three tails lack secondary and, therefore, tertiary structure (Fig. 3A). We next asked whether impaired processivity of Fhod-A and Fhod-E requires a specific region or simply relates to tail length. Because the tails are unfolded, we reasoned that we could truncate the tail without disrupting the formin structure. We truncated the last 24 residues of Fhod-A (Fhod-AΔ24, identical to truncating the last 18 residues of Fhod-E), the last 50 residues of Fhod-A (Fhod-AΔ50), and the last 75 residues of Fhod-A (Fhod-AΔ75). Truncating the last 99 residues of Fhod-A results in Fhod-B. We then analyzed actin elongation in the presence of each Fhod-A tail truncation via TIRF microscopy and compared these results to both Fhod-A and Fhod-B (Fig. 3C). All tested Fhod-A tail truncations processively elongated actin filaments based on the presence of dim filaments and their increased elongation rate relative to the bright filaments (Fig. 3C-D). The median run length (at t = 5 min) for each construct appears slightly shorter than that of Fhod-B (19.4 - 20.6 μm, number of filaments for each given in Fig. 3E), but there was no correlation between the decrease in processivity and the number of residues truncated (Fig. 3E). Together, these data show that Fhod processivity does not reflect the length of the tail and instead suggest that the last 24 residues of the Fhod-A tail (or the last 18 residues of the Fhod-E tail) are required to impair Fhod’s processivity (Fig. 3B).

**Figure 3.**
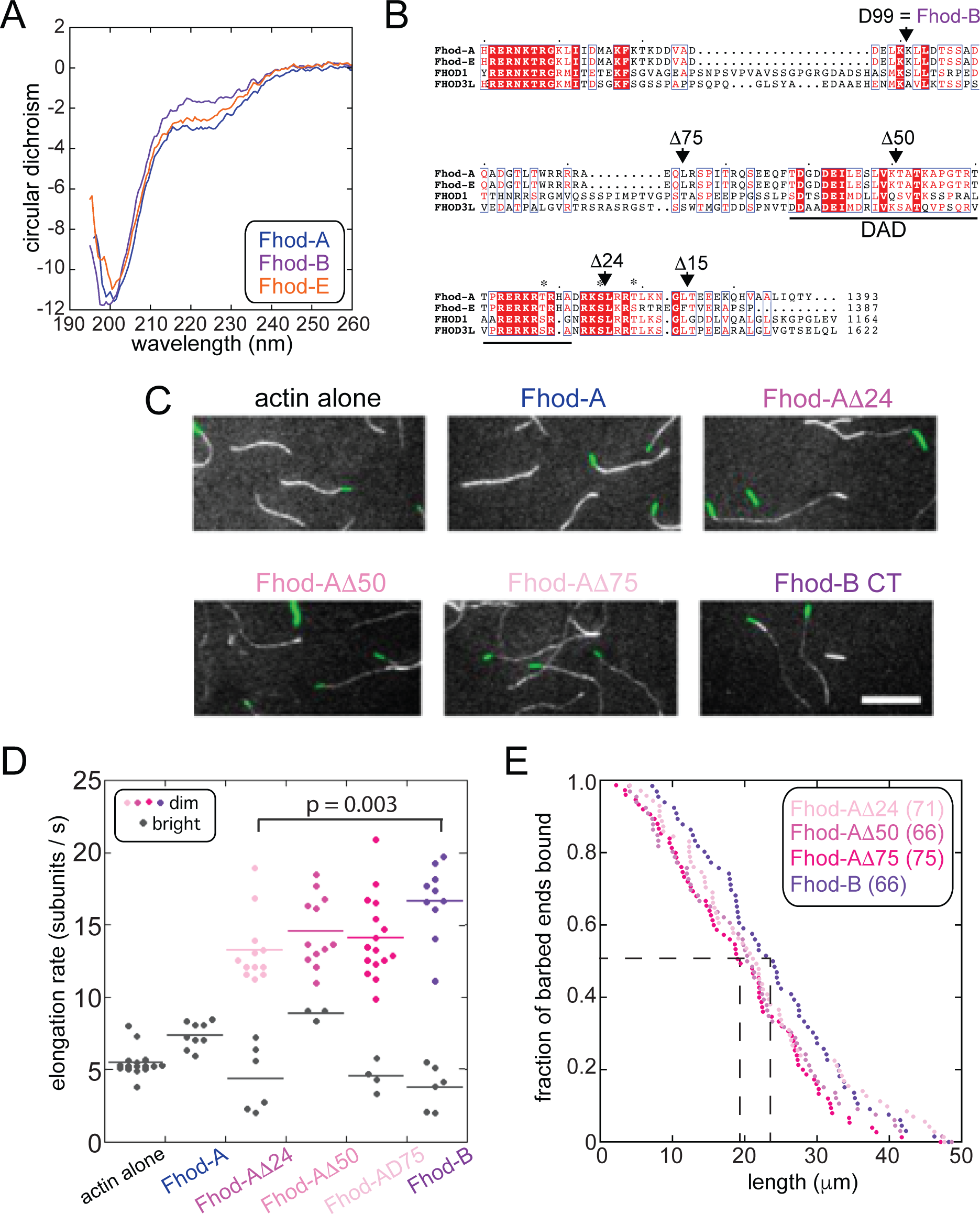
Residues at the end of the Fhod-A tail inhibit processivity. A) Wavelength scans of circular dichroism indicate that Fhod tails are disordered. B) Alignment of two Drosophila and two human Fhod tails. Truncation points for subsequent experiments are indicated. Asterisks denote phosphorylated residues (24). C) Direct observation of barbed-end elongation by TIRF microscopy. Conditions are the same as in figure 2. Images were taken 10 min after the start of polymerization. *Scale bar,* 10 μm. D) Quantification of elongation rates ≥ 2 flow chambers for each condition. Bright filaments are represented by grey dots and dim filaments are colored. Lines indicate the means. E) Measurement of processivity shown as an empirical CDF plot. Data points are from three experimental replicates (the number of filaments analyzed for each construct is given in the figure). The dashed lines indicates the medians. The mean +/-standard deviation of the three independent samples for each construct is 22.5 +/-0.3 μm (Fhodτι24), 20.8 +/-0.7 μm (Fhodτι50), 19.9 +/-0.6 μm (Fhodτι75), 24.1 +/-0.6 μm (Fhodτι99 = FhodB).

Examination of the primary sequences reveals a highly conserved basic region that extends beyond the traditional definition of the DAD but could be a continuation of this domain. This basic region straddles the Δ24 truncation site. To determine whether this region was responsible for loss of processivity, we made one further truncation, only removing 15 residues from the Fhod-A tail. Similar to wild-type Fhod-A, filaments grown in the presence of Fhod-AΔ15 were exclusively bright and grew at a rate indistinguishable from actin alone. The data indicate that the nine residues immediately adjacent to the DAD basic region are necessary to inhibit processivity, although we do not know if they are also sufficient.

We previously hypothesized that the Capu-tail increased processivity through electrostatic interactions with the actin filament. The pI’s of the Fhod-A and Fhod-E tails are higher than that of Fhod-B yet lower than or similar to other tails that support processivity (e.g. Capu and FMNL1 (13)) (Fig. 1B). Combined with the lack of correlation between processivity and length of truncation, we can rule out simple electrostatics to explain why Fhod-A and Fhod-E are not processive. We attempted to perform cosedimentation assays with isolated tails to measure their affinities for actin filaments. Unfortunately, due to non-specific binding at higher tail concentrations, the data did not plateau, and we could not make interpretable measurements.

### Fhod tails decrease nucleation and elongation

Formin tails also contribute to nucleation (12, 13). We, therefore, examined the impact of the different Fhod tails on the initial step of actin assembly. We performed both pyrene assays and TIRF microscopy. All three isoforms accelerated actin assembly compared to actin alone but at different rates (Fig. 4A). Assuming that elongation rates of each isoform are the same in the absence of profilin, the two isoforms with longer tails, Fhod-A and Fhod-E, are weaker nucleators than Fhod-B. We also spotted on coverslips actin polymerized in the absence or presence of Fhod isoforms. All three isoforms greatly enhanced the number of filaments per field of view (Fhod-E shown as representative), relative to actin alone, confirming that they promote actin assembly by nucleation (Fig. 4B,B’) (19).

**Figure 4.**
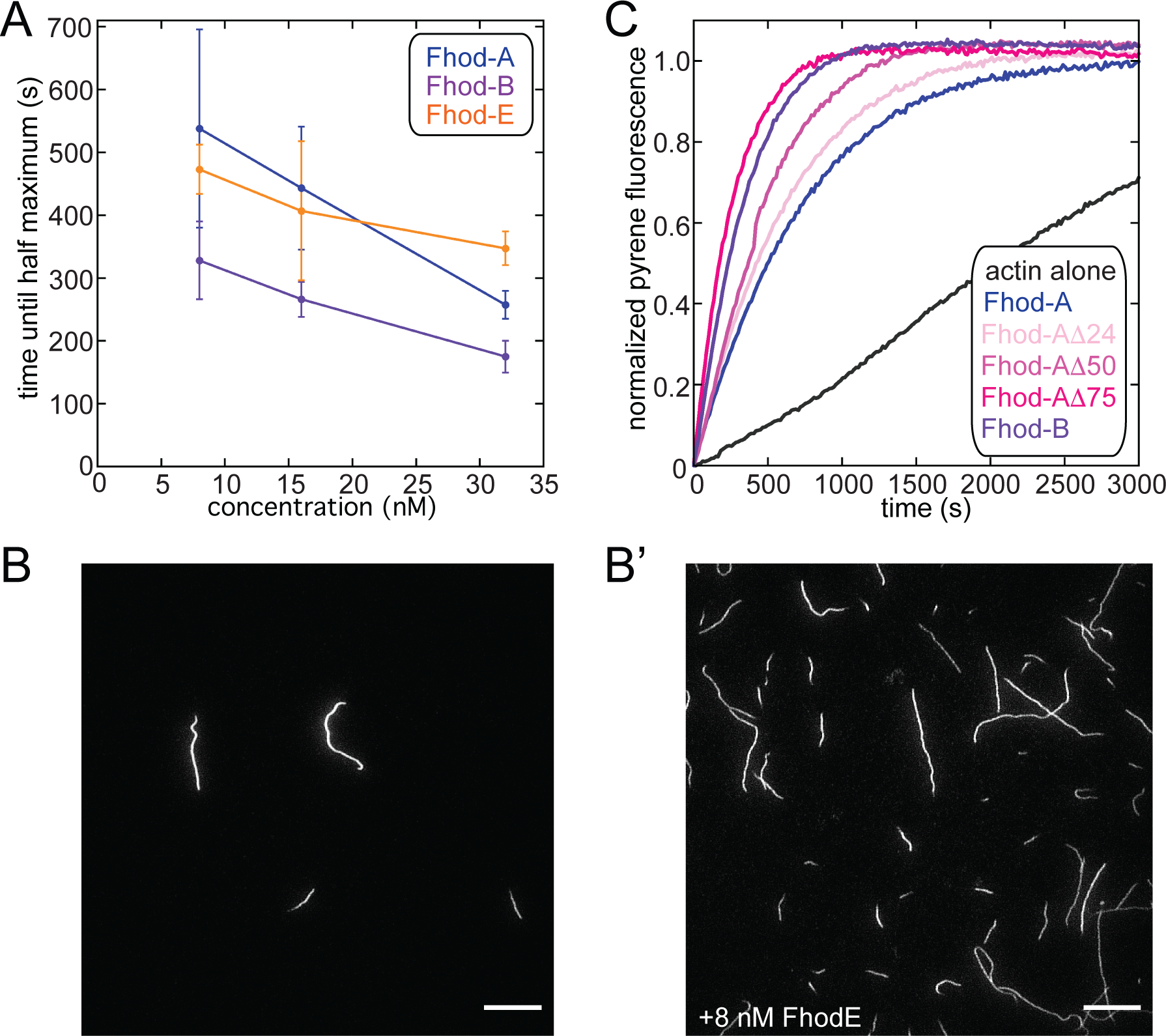
Fhod tails decrease nucleation. A) Assembly of 2 μM (10% pyrene-labeled) actin with 8-32 nM of indicated Fhod constructs. B) 2 μM actin was polymerized in the presence (B’) or absence (B) of 8 nM Fhod-E for 5 min, stabilized with Alexa Fluor 488-phalloidin, and imaged by TIRF microscopy. *Scale bars,* 10 μm.

We also measured the activity of the Fhod-A tail truncation constructs in pyrene assays. These assays suggest a trend with nucleation strength increasing as the tail is shortened (Fig. 4C). Similarly, the elongation rate increases gradually as the tail is shortened (Fig. 3D). The total change is small (13 ± 2 vs 17 ± 2 subunits/s; mean ± standard deviation, n = 12, 10) but statistically significant (p = 0.003; two tailed t-test with equal variance). Thus, the longer Fhod tails inhibit both nucleation and elongation, analogous to their effect on processivity but through a distinct mechanism correlating with tail length rather than depending on a specific region within the tail. Electrostatics may be important here. Fhod tail inhibitory activity contrasts with the tails of mDia1 and Capu, which increase nucleation but do not affect elongation (12, 13).

### Tails are major determinants of formin processivity

To examine how potent the enhancing and inhibitory effects of tails can be, we built chimeras (Fig. 5A). We chose Capu, a highly processive formin (λ ∼250 µm; Fig. 5D) and Delphilin, a formin that has little, if any, tail or processivity (Fig. 5B,C) (13, 20, 21). First, we asked whether the Fhod-A tail is sufficient to inhibit even the strong processivity of Capu. Indeed, when we replaced the 30-residue tail of Capu with the ∼100 residue tail of Fhod-A, all evidence of processivity was lost, and actin elongation was indistinguishable from actin alone (Fig. 5E). Because we previously found that the Capu construct lacking its tail (Capu1-1031) retains processivity (λ ∼75 µm) (13), we conclude that the Fhod-A tail inhibits processivity. To ask if a tail is sufficient to impart processivity, we added the tails of Fhod-B and Capu to the FH1FH2 of Delphilin (hDelFF). With the Fhod-B tail we observed only short dim stretches, not obviously different from those we occasionally observed for wild type Delphilin (Fig. 5C,F). On the other hand, the Capu tail greatly enhanced processivity of Delphilin, based on the long, dim filaments observed in these assays (Fig. 5D,G). Thus, formin tails can enhance the processivity of the FH2 domain, with the Capu tail even overcoming the inherently weak processivity of Delphilin. In sum, the tails are determining whether or not a formin is processive.

**Figure 5.**
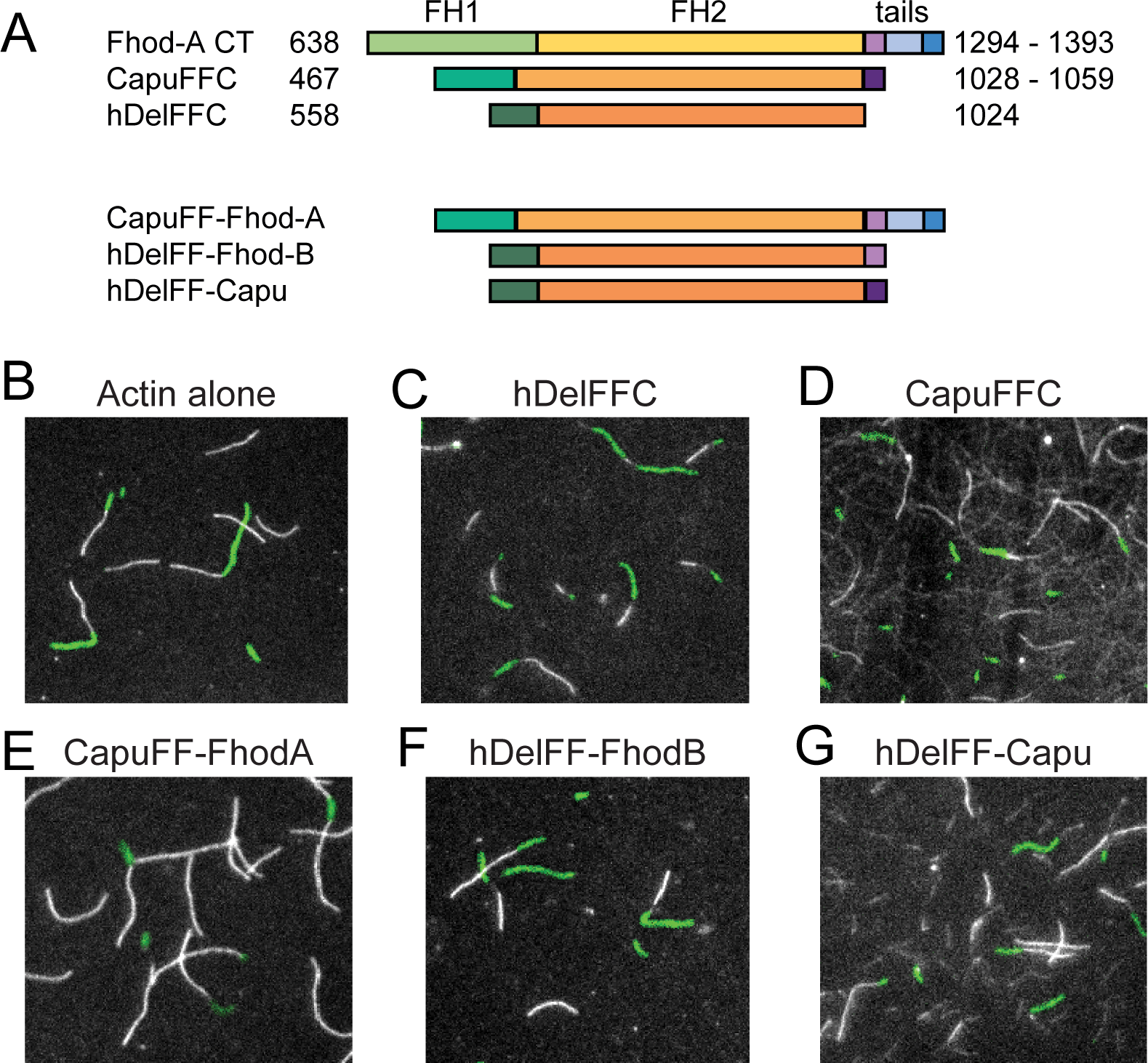
Tails dominate processivity. A) Cartoons of chimeras between FH1FH2 domains and tails of multiple formins. B) Direct observation of barbed-end elongation by TIRF microscopy. Conditions are the same as in figure 2 except, the indicated construct was added at different concentrations (hDelFFC = 15 nM, CapuCT = 1 nM, CapuFhodA = 0.5 nM, hDelFhodB = 15 nM, hDelCapu = 15 nM).

## Discussion

### Multiple Fhod tails are used

*Drosophila* Fhod contributes to many different functions, such as programmed cell death in the salivary gland, myogenesis, and immune response (15, 16, 18). *Drosophila* only have one Fhod gene, *fhos*, but nine splice variants are annotated in Flybase (14). We propose that splice variants are necessary for the wide range of roles. There is already clear evidence for differential expression of splice variants that vary in their N-termini: Fhod-A is sufficient to rescue the lethality and wing phenotypes of *Fhos*^Δ1^, a presumptive null (16). However, severe flight muscle phenotypes are observed when Fhod-A is the only variant expressed. Instead, Fhod-H, I, and/or J, are responsible for proper localization of Fhod and thin filament formation (15). These longer isoforms contain distinct 5’ regions, including exons and introns, but they all have the same tail as Fhod-A. While the N-terminal half of formins contribute to autoinhibition it is also the most variable region and is often responsible for specific localization and/or binding interactions.

We asked if by changing the C-terminus of the formin, one could alter the actin assembly activity, essentially creating formins capable of building distinct structures. Our data indicating that Fhod-A tail mRNA is present through development is consistent with the known ability of Fhod-A to rescue viability. Interestingly, mRNA of the Fhod-E tail is expressed only after eclosion. The Fhod-E tail is expressed at increasing levels in the young adult, eventually reaching a level comparable to Fhod-A (based on read number), suggesting that it plays a widespread role in the adult fly. In contrast, Fhod-B and -D were not detectable by this method. Each represents a unique transcript. In preliminary RT-PCR experiments, we saw evidence of both Fhod-D and -B but at levels too low to quantify (data not shown). Perhaps they are each expressed in only one or a few tissue types. In fact, the Fhod-B reading was so low that we question whether or not it is translated in the cell. Notably, Fhod-B lacks a DAD domain and, presumably, cannot be autoinhibited. If this is truly a constitutively active formin, it seems judicious to tightly control its expression levels.

### Processivity

Processivity (the dissociation rate of a formin from a barbed end) and the elongation rate determines the average filament length built by a formin (also called the characteristic run length). Studies of processivity show that the dissociation rate is correlated with the elongation rate, suggesting that the FH2-barbed end interaction is weakened in response to addition of actin monomers to the barbed end (9, 22). In addition, it has been shown that mDia1’s processivity is highly sensitive to force (22). Processivity is presumably a property of the specific FH2 domain bound to a barbed end. Experiments with altered ionic strength show that electrostatic interactions play an important role, and molecular dynamics simulations of bound barbed ends provide corroborating evidence for electrostatic interactions at the FH2-barbed end interface (22, 23). However, processivity is also influenced by the two regions that straddle the FH2 domain: the FH1 domain and the C-terminal tail (13, 22). Processivity enhancement by the FH1 domain depends on profilin and is proposed to be stabilization by ring closure, suggesting that the weak state, perhaps translocation of the trailing FH2 half, occurs upon actin monomer addition. Enhancement by the tail was proposed to be through non-specific electrostatic interaction between the tails and the actin filament, which aligns with ionic strength assays (13, 22). Perhaps not surprisingly, we found that determination of processivity is more complex in some cases.

At first, we did not know if the low processivity of Fhod-A (λ = ∼2 um) was inherent to the FH2 domain and/or due to its tail. Based on our previous observations with Capu, and the length of the FhodA-tail (99 AA), we hypothesized that the FH2 domain was not capable of processive elongation and even a long tail could not overcome the weak interaction. Instead, we found that Fhod-B, with a shorter tail (that is included in the Fhod-A tail), is ten-fold more processive. It follows that the Fhod-FH2 domain can support processive actin elongation but the time that it remains bound depends on the tail. Consistent with this conclusion, we found that the tail determined whether or not multiple chimeras were processive. Notably, adding the Capu tail to Delphilin enabled Delphilin to processively elongate filaments.

There appears to be a marked difference between Fhod and Capu in how their FH2 domains and tails contribute to processivity. The data demonstrate that tail length and pI, which correlate with processivity in some formins, do not predict processivity for Fhod isoforms. So how does the Fhod-A tail inhibit processivity of the FH2 domain? We found that a stretch of basic amino acids extending beyond the DAD is necessary to inhibit processivity. All three long Fhod tails as well as both mammalian Fhod isoforms contain additional basic residues that extend past the typical DAD. This basic region might directly bind to the FH2 domain to decrease its processivity. We do not favor this model because the Fhod-A tail was also able to inhibit processivity by the unrelated FH2 domain of Capu. Alternatively, the extended basic region could bind the sides of actin filaments “too tightly”, pulling the FH2 off of the barbed end instead of loosely stabilizing it. As stated, we were unable to test this model directly.

Within the DAD domain and the adjacent inhibitory region that we identified are three well-documented phosphorylation sites (Fig. 3B). Studies with both Fhod-A and mammalian homologs show that Rho-dependent kinase (ROK/ROCK) phosphorylates these sites (16, 24). Phosphorylation of these residues is sufficient to release the autoinhibitory interaction in pull down assays and activate the formins in tissue culture experiments (16, 24). For example, expressing Fhod3 with phosphomimetic S -> D mutations in HeLa cells resulted in higher filamentous actin levels compared to expressing wild-type Fhod. However, the analogous mutations did not alter the chemistry of the tail enough to make Fhod-A processive (not shown). This leaves us with the question of whether there is a mechanism to convert Fhod-A and -E into more processive elongation factors. Alternatively, low processivity may be essential to building shorter filaments, such as the thin filaments in sarcomeres and stress fibers that are assembled by Fhod-family formins. Thin filaments are typically ∼1 μm long. Thus, Fhod-A’s characteristic run length of ∼2 μm is sufficient and well-suited to build such structures. Perhaps, Fhod-family formins have evolved away from the highly processive formins.

In sum, the role of formins tails was originally believed to be restricted to all of nothing regulation through autoinhibition. In fact, many experiments have been performed with constructs lacking the tail, in order to create a constitutively active formin. We now know that formin tails can have dramatic effects on nucleation and elongation and, therefore, should not be ignored.

## Methods

### Construct cloning and purification

All constructs were built from Fhod isoform A cDNA (clone SD08909, obtained from the *Drosophila* Genomics Resource Center). The cDNA was used as a template to clone C-terminal constructs into either a modified version of the pET-15b plasmid with an N-terminal His_6_ tag (Fhod-A, Fhod-AΔ24, Fhod-AΔ50, Fhod-AΔ75, and Fhod-E) or a pGEX-6P-2 plasmid (Fhod-B). All truncated tail constructs were produced via FastCloning (25). The Fhod-E exon was synthesized and added by Gibson Assembly.

#### Fhod-A, Fhod-AΔ24, Fhod-AΔ50, Fhod-AΔ75, and Fhod-E

Purification was performed as described (19). In brief, these constructs were purified using a HitrapSP-FF cation exchange column (GE Life Sciences) followed by a MonoQ anion exchange column (GE Life Sciences). Peak fractions were dialyzed into Fhod storage buffer (10 mM Tris, pH 8.0, 150 mM NaCl, 1 mM DTT). The purified constructs were aliquoted, flash frozen using liquid nitrogen, and stored at −80°C.

#### Fhod-B

Purification was carried out using glutathione sepharose resin. The eluted protein was dialyzed in PBS with 1 mM DTT and cleaved with Precission Protease overnight. The cleaved protein was filtered through glutathione sepharose and dialyzed into 10 mM Tris, pH 8.0 50 mM NaCl, 1 mM DTT. It was next run on a MonoQ anion exchange column (GE Life Science). Peak fractions were dialyzed into Fhod storage buffer. Aliquots were stored at −80°C.

*Acanthamoeba castellanii* actin and *Drosophila* profilin (Chic) were purified as described previously (26, 27). Actin was labeled with pyrene-iodoacetamide (27), Oregon Green 488-iodoacetamide (Invitrogen) (26), or EZ-link maleimide-PEG2-biotin (Thermo Scientific) (28) as described.

### Pyrene assays

Bulk actin polymerization assays were performed on an Infinite 200 Pro plate reader (Tecan) essentially as described (26). In brief, 2 μM 10% pyrene-labeled actin monomers were incubated for 2 min in ME buffer (200 μM ethylene glycol tetraacetic acid [EGTA] and 50 μM MgCl_2_) to convert Ca^2+^-actin to Mg^2+^-actin. Polymerization was initiated by adding KMEH buffer (final concentration: 10 mM Na-HEPES, 1 mM EGTA, 50 mM KCl, and 1 mM MgCl_2_) to the Mg^2+^-actin. Fhod constructs were added to the KMEH buffer before addition to Mg^2+^-actin. To complement nucleation assays, we polymerized actin under the same conditions and stopped the reaction after 5 minutes by adding Alexa Fluor-488 phalloidin. Samples were then diluted and spotted on poly-L-lysine coated coverslips.

### TIRF microscopy

TIRF microscopy was utilized to measure the elongation rates and processivity of the Fhod constructs. Biotinylated coverslips were prepared as described (19). Flow chambers of ∼15 μl were assembled on slides with strips of double-sided tape. Flow chambers were prepared with the following steps: 1) incubated for 2 min with block containing 25 μl of 1% Pluronic F-127 (Sigma), 50 μg/ml casein in PBS; 2) washed with 25 μl of KMEH; 3) incubated for 1 min with 25 μl of 40 nM streptavidin in KMEH; 4) washed with 25 μl of TIRF buffer (KMEH, 0.5% methylcellulose (400 cP, Sigma), 50 mM DTT, 0.2 mM ATP, 20 mM glucose). Fhod was incubated with F-actin seeds 45 s prior to addition of Mg^2+^-G-actin. Oregon green-labeled G-actin with incubated with *Drosophila* profilin for 2 min in ME buffer to convert Ca^2+^-actin to Mg^2+^-actin. The final concentrations in the flow chambers were as follows: 1 μM Mg^2+^-G-actin (20% Oregon green labeled) in KMEH, 5 μM profilin, 0.5 nM (unless otherwise indicated) Fhod construct, 0.5 nM F-actin seeds (1% biotinylated, stabilized with Alexa Fluor 647-phalloidin), 250 μg/ml glucose oxidase, 50 μg/ml catalase, and 50 μg/ml casein in TIRF buffer. Most experimental data were acquired with a DMI6000 TIRF microscope (Leica, Germany) with an HCX PL APO objective (100× magnification, N.A. = 1.47), and an Andor DU-897 camera, using the Leica application suite advanced fluorescence software. The data in Fig. 5 were acquired with a Zeiss Axio Observer 7 Basic Marianas Microscope with Definite Focus 2 equipped with a 3i Vector TIRF System, an Alpha Plan-Apochromat 63x/1.46NA Oil TIRF Objective, and an Andor iXon3 897 512×512 10MHz EMCCD Camera, using Slidebook 6 software. Experiments were performed at room temperature. Images were captured at 10 s intervals for 10 min. Filament lengths were quantified with the JFilament plug-in in FIJI (29).

### Circular Dichroism Spectroscopy

Freshly purified protein stocks were dialyzed into 50 mM potassium phosphate (pH 7.5), 1 mM DTT and then precleared by centrifugation at 100,000 x g for 20 min at 4 °C. Circular dichroism (CD) spectra were measured on a J-715 spectropolarimeter (Jasco, Tokyo, Japan) by averaging two wavelength scans from 195 to 260 nm.

#### Nanopore Sequencing

*D. melanogaster* w^1118^ flies were collected at different developmental stages: third instar larva, pupa, and days 1, 3, and 5 post-eclosion. Total RNA was extracted by freezing flies for 10 minutes in TRIzol Reagent (ThermoFisher Scientific, catalog #15596026) and grinding with a pestle. Homogenized samples were then centrifuged at 12k rpm for 30 seconds to remove fly debris. The supernatant was transferred to a new tube, chloroform extracted, and centrifuged at 10k rpm for 15 minutes. The rest of the RNA purification was performed using the Direct-zol RNA MiniPrep Kit (Genesee Scientific, catalog #11-330) and Zymo-Spin IIICG columns (Zymo Research, catalog #C1006-50-G) as per the manufacturer’s instructions with a minor change in the flowthrough step, which was centrifuged for 2k rpm for 2 minutes and repeated once more. cDNA was synthesized using SuperScript III (ThermoFisher Scientific, catalog #18080093) using an Oxford Nanopore sequence 5’ ACTTGCCTGTCGCTCTATCTTC oligo(dT)_18_ 3’. Ethanol precipitation was performed to purify and concentrate the cDNA. The purity and concentration were assessed with a NanoDrop. The cDNA was then amplified with Phusion High-Fidelity DNA Polymerase (New England Biolabs, catalog #M0530S) using the following steps: initial denaturation 98℃, 0:30; 30 cycles 98℃, 0:30, 64℃, 0:30, 72℃, 1:30, final extension 72℃, 5:00, and infinite hold 4℃. Standard 40 uL reaction was followed but with a 3x increase in forward primer.

Forward primer: 5’ TTTCTGTTGGTGCTGATATTGCATGATACCGCAGGTGGTGGG 3’ (Note that the Nanopore sequence TTTCTGTTGGTGCTGATATTG is not needed after switching from the PCR Sequencing Kit to the Native Barcoding Kit but was used for ease).

Reverse primer: 5’ ACTTGCCTGTCGCTCTATCTTC 3’ Ethanol precipitation was performed to purify and concentrate the PCR amplicons. The purity and concentration were assessed using a NanoDrop.

Nanopore libraries were prepared from 130 ng of Fhos amplicons using the Native Barcoding Kit 24 V14 from Oxford Nanopore (ONT, catalog #: SQK-NBD114.24) as per the manufacturer’s instructions. Sequencing was performed using R10.4.1 flow cells on a MinION Mk1B device and sequenced for 48 hours. Basecalling was performed using Guppy Basecaller (Version 6.4.6). Reads were then mapped to a *FHOS* reference sequence obtained from NCBI (Gene ID: 39004) using Minimap 2 (Version 2.17-r941). Reads were visualized using IGV (Version 2.12.3) and figures were prepared using Inkscape (Version 1.1.2).

### Data availability

All data will be shared upon request sent to the corresponding author.

This article contains supporting information.

## Acknowledgements

The authors would like to thank the entire Quinlan lab for support and useful discussions throughout this project. Thanks especially Margo Myers and Bryan Christian, who did some important thinking and supporting experiments. Thanks also to Theodosia Bartaschevitch for qPCR experiments that inspired the nanopore sequencing work.

## Funding

This work was supported by National Institutes of Health Grants under Award Numbers T32GM145388 (to C.W.), R35 GM130370 (to G.C), and R01 HL14659 (to M. E. Q.). The content is solely the responsibility of the authors and does not necessarily represent the official views of the National Institutes of Health.

The authors declare that they have no conflicts of interest with the contents of this article.

---

**Figure S1.**
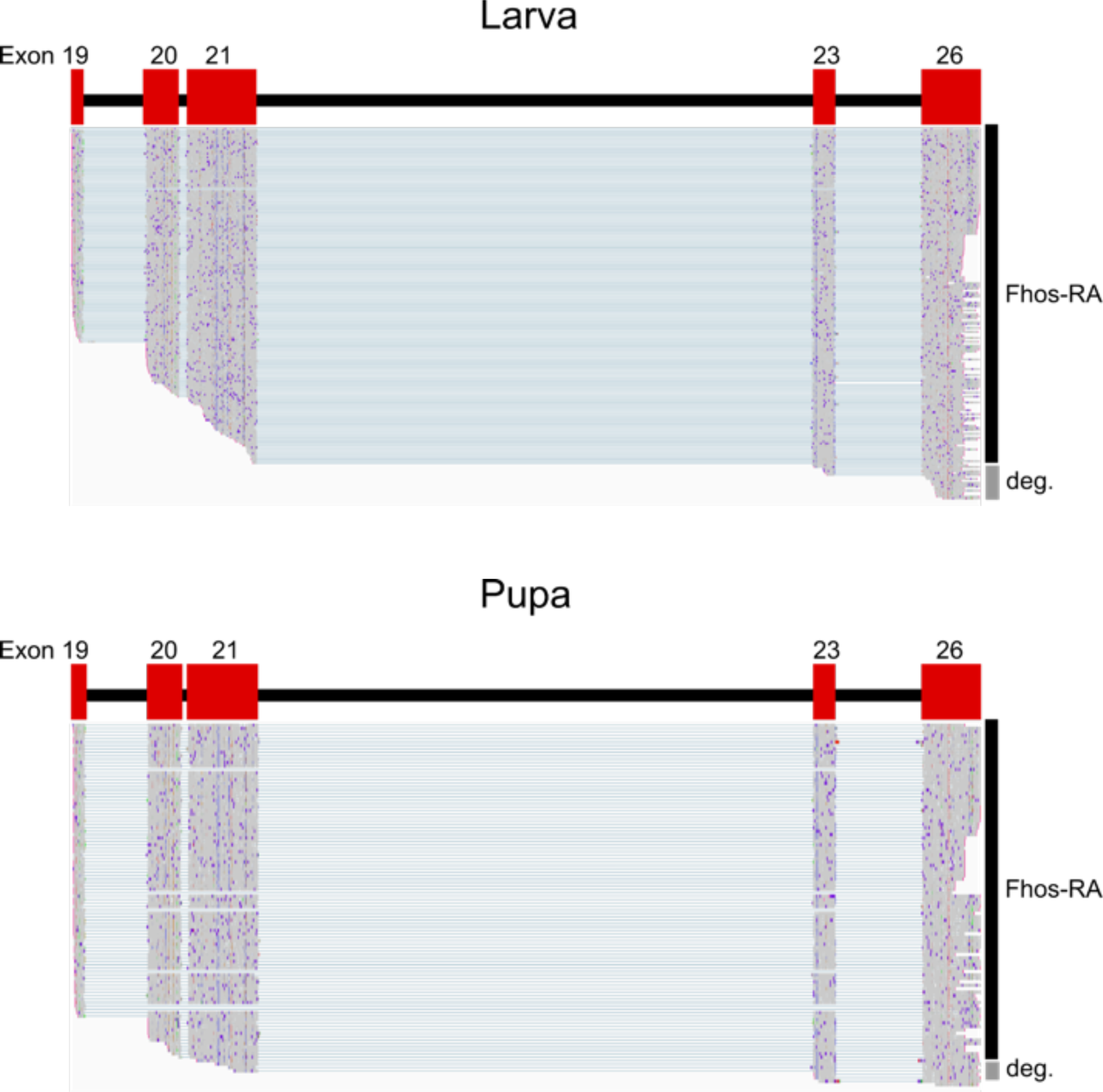
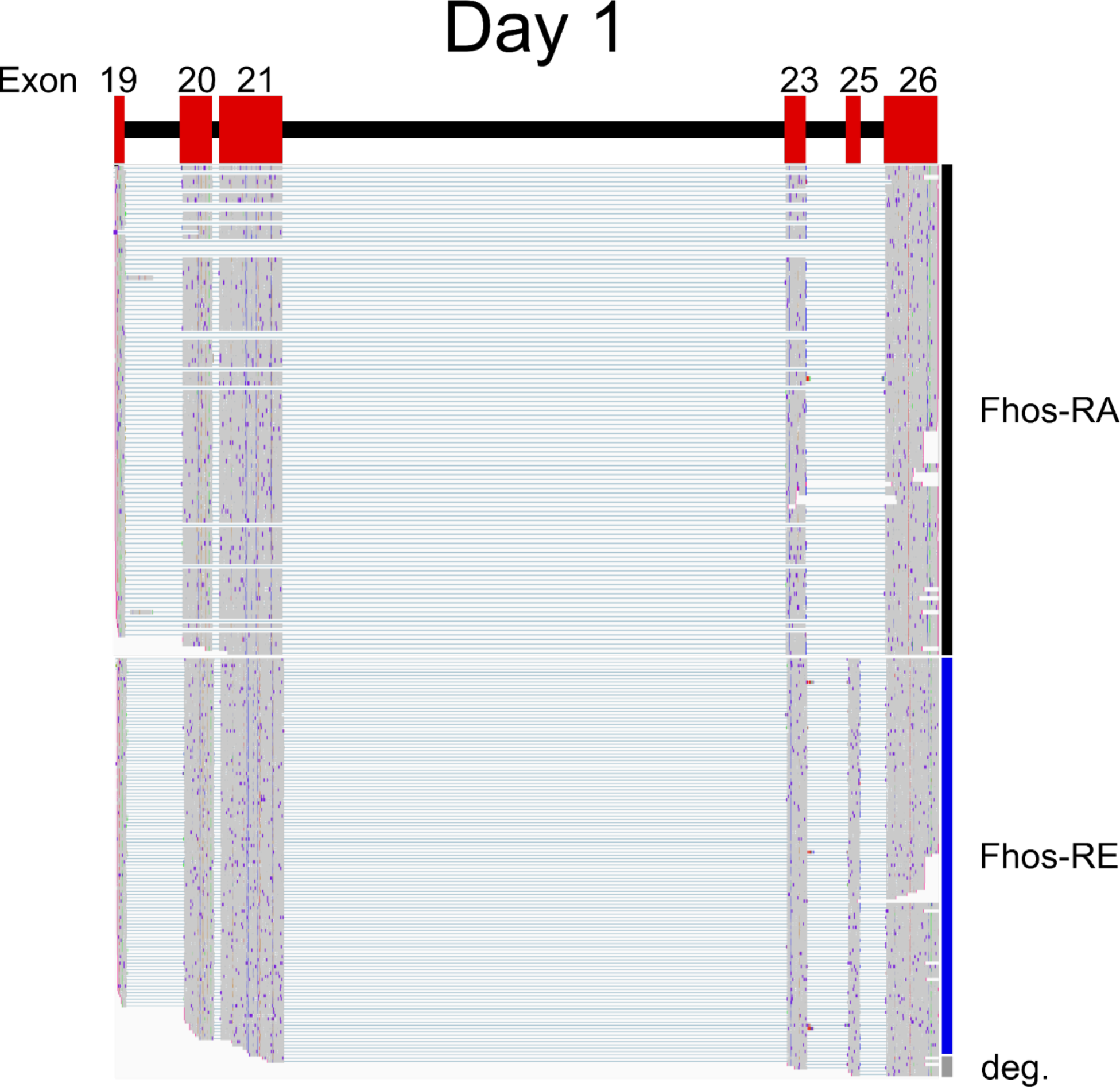
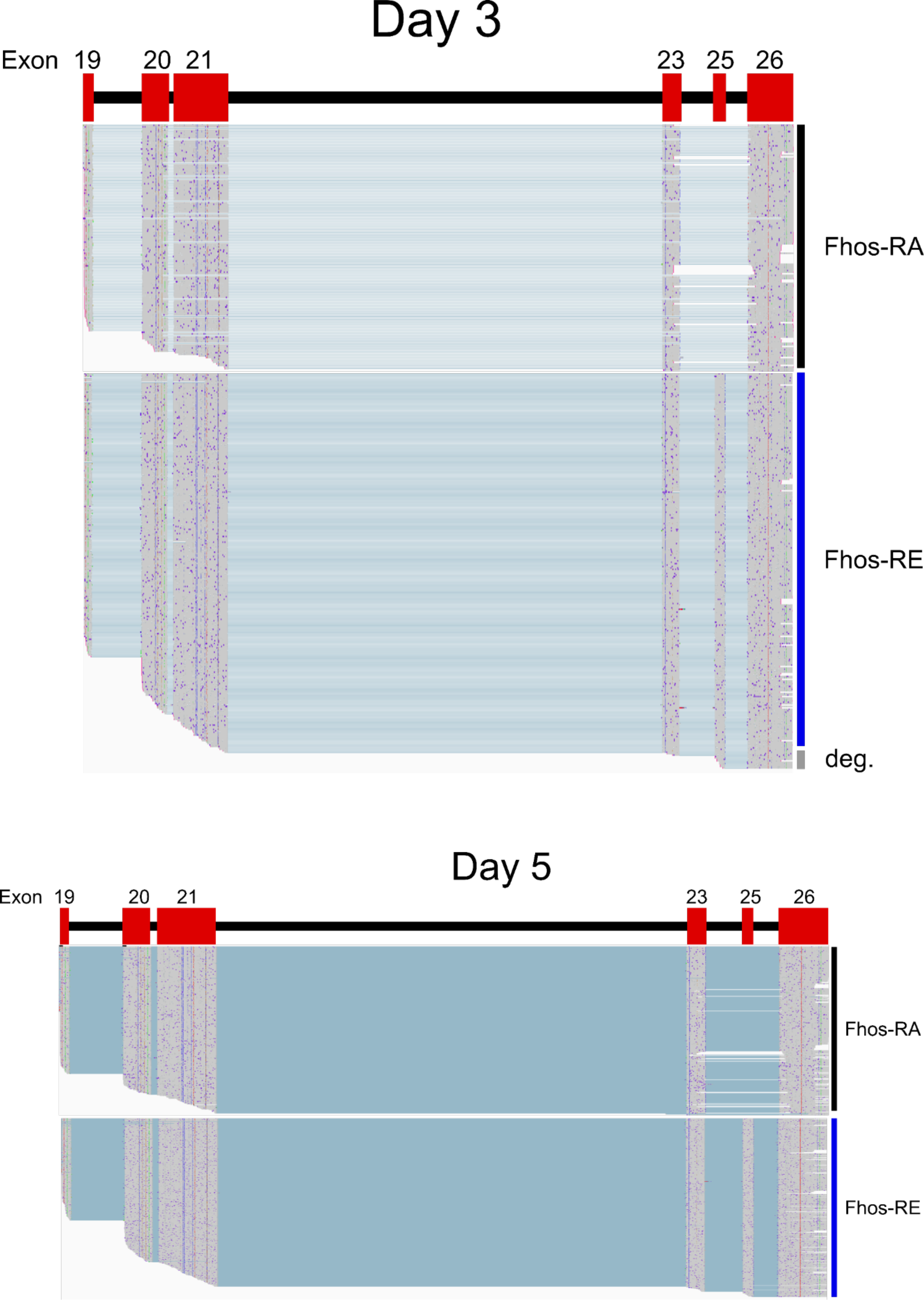
Oxford Nanopore sequencing – samples of reads mapped to the Fhod gene for the five developmental stages assessed.

**Figure S2.**
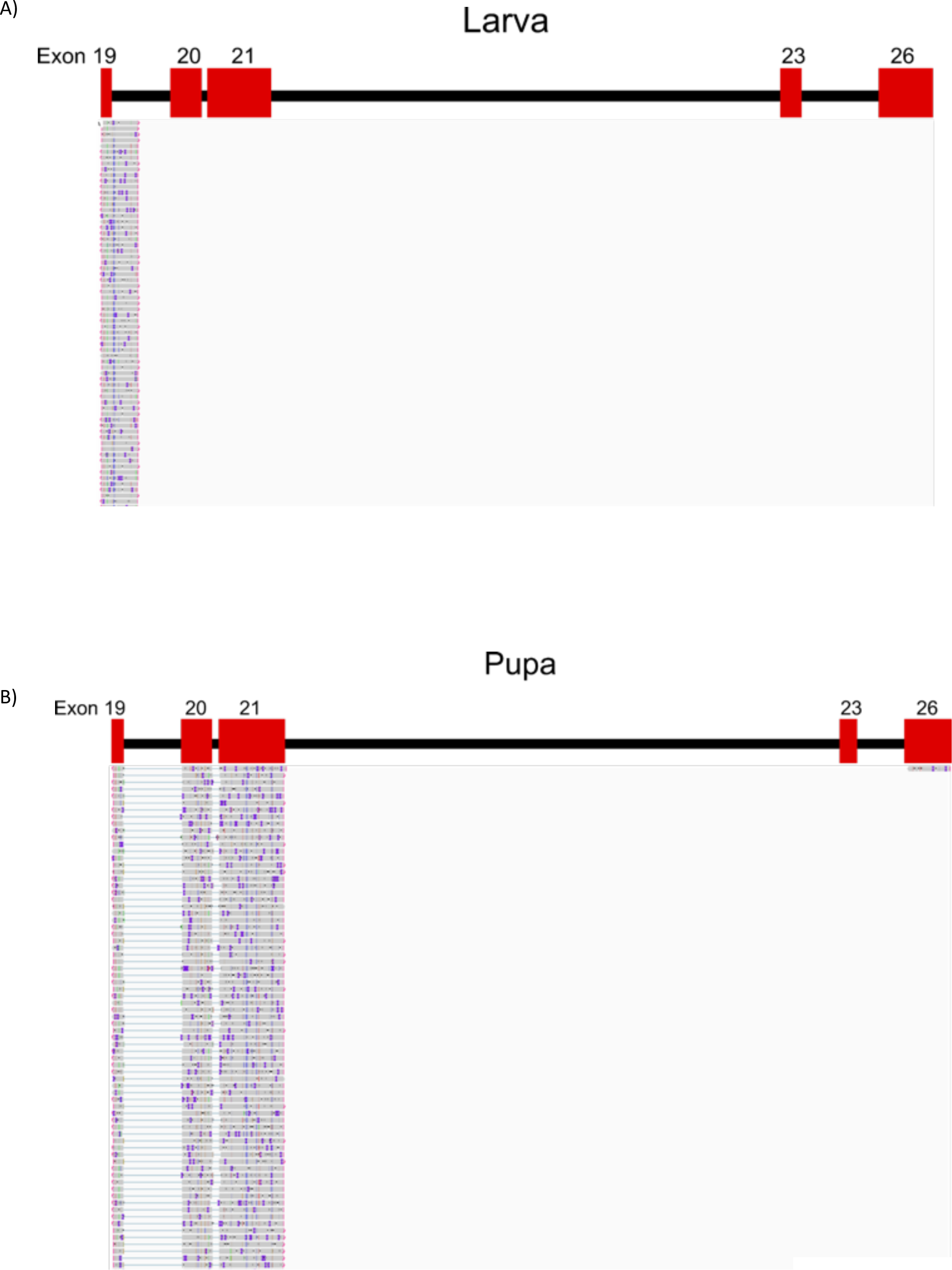
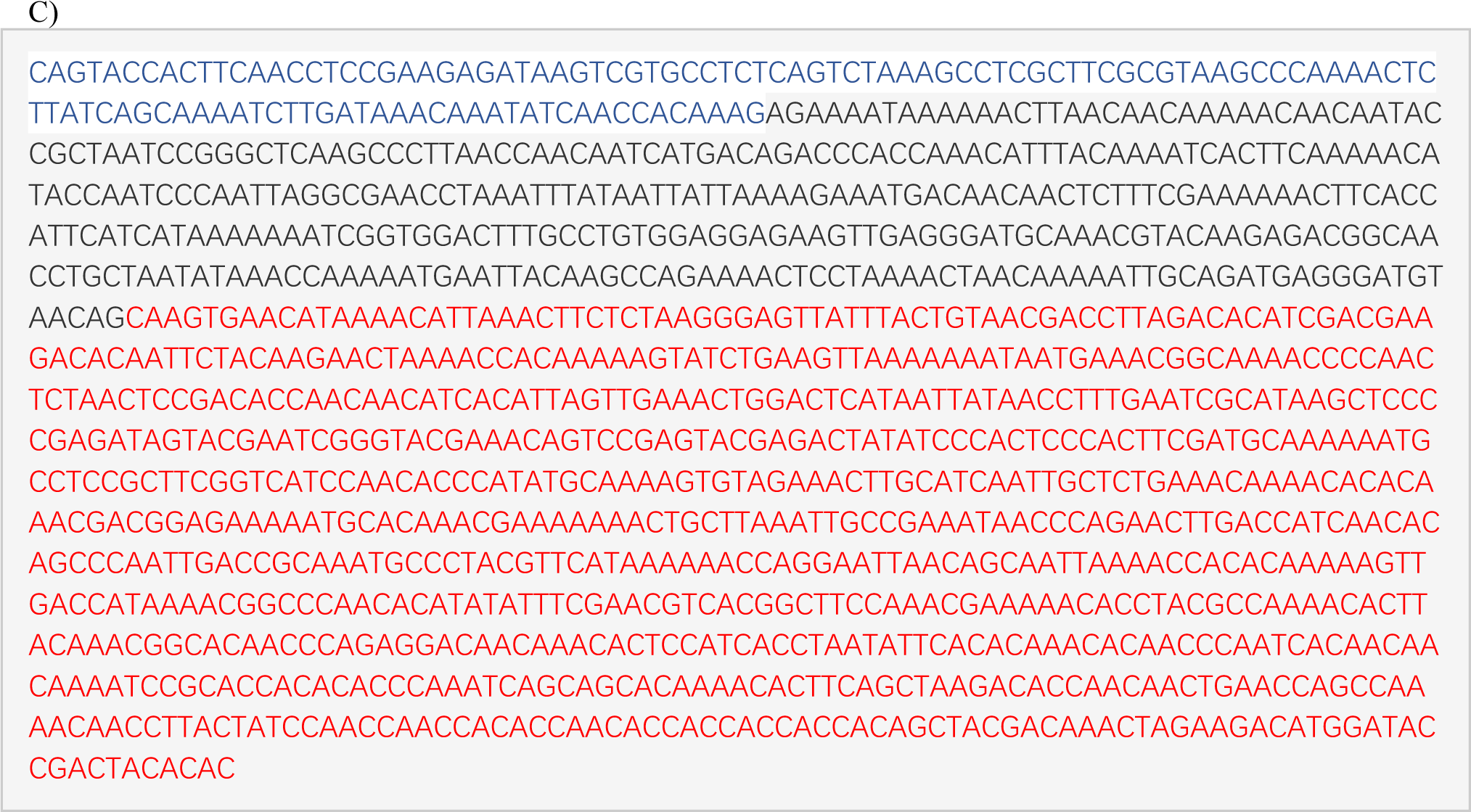
Fhod variants that accumulated, which have not been previously annotated. A) Detection of reads that show the extended region of exon 19. B) Detection of reads that have only exons 19, 20, and 21. C) Sequence of the few reads that detected only the distinct Fhod-B 3’ UTR + extended region. Blue = Fhod-B UTR, black = extended Fhod-B intronic region matching FlyBase ID: FBcl0104834, red = continuation of intronic region.

## Notes

### Competing Interest Statement

The authors have declared no competing interest.

